# Construction of a potent pan-vaccine based on the evolutionary tendency of SARS-CoV-2 spike protein

**DOI:** 10.1101/2021.12.21.473594

**Authors:** Yongliang Zhao, Wenjia Ni, Simeng Liang, Lianghui Dong, Min Xiang, Zeng Cai, Danping Niu, Qiuhan Zhang, Dehe Wang, Yucheng Zheng, Zhen Zhang, Dan Zhou, Wenhua Guo, Yongbing Pan, Xiaoli Wu, Yimin Yang, Zhaofei Jing, Yongzhong Jiang, SARS-CoV-2 Vaccine Task Force Group, Yu Chen, Huan Yan, Yu Zhou, Ke Xu, Ke Lan

**Author notes:** Corresponding authors. Address correspondence and reprint requests to Dr. Ke Lan) and Dr. Ke Xu). These authors contributed equally to this work (Yongliang Zhao, Wenjia Ni, Simeng Liang, Lianghui Dong, Min Xiang, Zeng Cai, Danping Niu).

## Abstract

SARS-CoV-2 continued to spread globally along with different variants. Here, we systemically analyzed viral infectivity and immune-resistance of SARS-CoV-2 variants to explore the underlying rationale of viral mutagenesis. We found that the Beta variant harbors both high infectivity and strong immune resistance, while the Delta variant is the most infectious with only a mild immune-escape ability. Remarkably, the Omicron variant is even more immune-resistant than the Beta variant, but its infectivity increases only in Vero E6 cells implying a probable preference for the endocytic pathway. A comprehensive analysis revealed that SARS-CoV-2 spike protein evolved into distinct evolutionary paths of either high infectivity plus low immune resistance or low infectivity plus high immune resistance, resulting in a narrow spectrum of the current single-strain vaccine. In light of these findings and the phylogenetic analysis of 2674 SARS-CoV-2 S-protein sequences, we generated a consensus antigen (S_6_) taking the most frequent mutations as a pan-vaccine against heterogeneous variants. As compared to the ancestry SWT vaccine with significantly declined neutralizations to emerging variants, the S_6_ vaccine elicits broadly neutralizing antibodies and full protections to a wide range of variants. Our work highlights the importance and feasibility of a universal vaccine strategy to fight against antigen drift of SARS-CoV-2.

## Introduction

Since the outbreak of the SARS-CoV-2 in late 2019, it has caused a global pandemic that is still going on (*1–3*). Globally, on December 14, 2021, there have been 270,031,622 confirmed cases of COVID-19, including 5,310,502 deaths, reported to WHO (https://covid19.who.int/). As an RNA virus of long genome size, the SARS-CoV-2 has a relatively high mutation frequency so that various mutations have appeared after nearly two years of continuous transmission (https://bigd.big.ac.cn/ncov/variation/annotation), especially mutations on S protein would change the infectivity and antigenicity of the virus (*4*). The firstly identified prevalent mutation was D614G located in the SD2 domain, which altered the fusion-competent confirmation of S protein and enhanced the ACE2-binding ability (*5*). Subsequently, the A222V mutation detected in Spain spread quickly throughout Europe in the middle of 2020 (*6*). As the different mutants occur constantly, the term “variant” has been used to denote viral strains that carry a series of co-emerging amino-acid mutations (*7*). The WHO has recently announced a Greek alphabet-based naming scheme for variants, including variants of concern (VOC) and variants of interest (VOI) (https://www.who.int/activities/tracking-SARS-CoV-2-variants/tracking-SARS-CoV-2-variants).

The VOCs are Alpha (B.1.1.7, first detected in the UK) (*8*), Beta (B.1.351, first detected in South Africa) (*9*), Gamma (P.1, first detected in Brazil) (*10*), Delta (B.1.617.2, first detected in India) (*11*), and Omicron (B. 1.1.529, first detected in South Africa) (*12*). The VOIs are Eta (B.1.525) (*13*), Kappa (B.1.617.1 first detected in India) (*11*), and Lambda (C.37 first detected in Peru) (*14*). The Alpha variant (B.1.1.7) carried 9 coordinated mutations including 2 deletions of amino acids at positions 69-70 (del69-70) and 144 (del144) of the NTD in S protein, N501Y/A570D located in the RBD, D614G/P681H located in the SD2, T716I/S982A located in the HR1, and D1118H (*8*). The Beta variant (B.1.351) contained 10 coordinated mutations that are L18F/D80A/D215G/R246I and amino-acids deletion at position 242-244 (del242-244) located in the NTD, K417N/E484K/N501Y located in the RBD, D614G located in the SD2, and A701V (*9*). The Gamma variant (P.1) possessed 11 coordinated mutations that are L18F/T20N/P26S/D135Y/R190S located in the NTD, K417T/E484K/D614G/N501Y/H655Y located in the SD2, and T1027I (*10*). The Delta variant (B. 1.617.2) was earliest documented in India in October 2020, which had 6 coordinated mutations including T19R located in the NTD, L452R/T478K located in the RBD, D614G/P681R located in the SD2, and D950N located in the HR1 (*15*). The Omicron variant (B. 1.1.529) contains more mutations (~32-35 mutations) than previous variants in which 15 mutations are located in the RBD (*12*).

With the emergence and widespread popularity of various variants, most likely viral evolution possesses a certain rationale behind it. The Alpha variant (B.1.1.7) was documented in December 2020 and more than 80% of globally new infectors have been identified to be the Alpha variant between April and May 2021 (https://cov-lineages.org/global_report_B.1.1.7.html). The Beta variant (B.1.351) was detected after the Alpha variant (B.1.1.7), the first clinical sample was documented on 8 October 2020, and the Beta variant (B.1.351) became the dominant variant in South Africa within one month (Tracking the international spread of SARS-CoV-2 variants B.1.1.7 and B.1.351/501Y-V2 (https://virological.org/t/tracking-the-international-spread-of-sars-cov-2-lineages-b-1-1-7-and-b-1-351-501y-v2/592). Epidemiological studies and mathematical modeling suggested that the RBD mutation N501Y increased transmissibility of up to 90% above other variants and results in higher nasopharyngeal viral loads than the original variant (*16*). It was also reported that the mutations of the NTD facilitated the Beta variant (B.1.351) to escape the mAb therapies or vaccines (*17, 18*).The E484K and N501Y mutations in the RBD contributed to neutralizing antibodies resistance (*19*). The Delta variant (B.1.617.2) was identified in October 2020 in India and spread faster than other variants (60% more transmissible than the Alpha variant) (*11*). To date, over 82% of global new infectors were identified as the Delta variant (B. 1.671.2) (https://covid.cdc.gov/covid-data-tracker/#variant-proportions). Recent studies indicated that the L452R mutation could lead to increased viral load in patients (*20*). However, the regularity underlying all these mutations is still unclear.

To curb the spread of the COVID-19 pandemic, different types of vaccines have been licensed but most of them were still based on ancestral WT spike protein. It has been reported that the sera of individuals injected one or two doses of the ancestral mRNA vaccines, either Moderna (mRNA-1273) or Pfizer/BioNTech (BNT162b2), had a decreased neutralization against the Alpha variant (B.1.1.7), Gamma variant (P.1) and Kappa variant (B.1.617.1) (about 2-3 fold, 2-7 fold or 6-7 fold respectively), and sharply decreased neutralization against Beta variant (B.1.351) (about 27.7-34.5 fold) (*21, 22*) wherein even more severe decrease occurred to Omicron variant (B.1.1.529) (about 10-41.1 fold) (*23, 24*). The sera of individuals who had been injected one or two doses of AstraZeneca-Oxford-ChAdOx1 also showed decreased neutralization against the Alpha variant (B.1.17) (~2.1 fold) and the Beta variant (B.1.351) (~9 fold) (*25*). The over-spread of the Delta variant (B.1.617) and the recent Omicron variant (B.1.1.529) have also raised concerns about the effectiveness of the vaccines approved so far. Therefore, redesign and update of SARS-CoV-2 vaccines to be universally protective to heterogeneous variants is in urgent need.

In the present study, we defined the infectivity and immune-escape ability of 54 (covering 45 single mutations, 8 variants, and WT) SARS-CoV-2 pseudoviruses, and then analyzed the evolutional route of SARS-CoV-2 spike protein. We found that SARS-CoV-2 spike protein evolves into distinct evolutionary paths of either high infectivity plus low immune resistance or low infectivity plus high immune resistance, resulting in heterogenous antigenicity. Thus, the current single-strain vaccine could provide narrow-spectrum protection only to one group of variants but not to the others. By phylogenetic analysis of 2674 S proteins, we generated a consensus S_6_ vaccine taking the most frequent mutations that could elicit broadly strong neutralization to all the VOC and VOI variants up to date. This study light with new hope that a universal SARS-CoV-2 vaccine is feasible and superior when facing continuous viral mutagenesis.

## Results

### The infectivity of SARS-CoV-2 variants

The frequencies of SARS-CoV-2 variants from the beginning of the COVID-19 pandemic until Dec 2021 were shown in **Figure 1a**. The number of ancestral WT viruses decreased dramatically from March 2020 and almost vanished until Oct 2020 wherein the Alpha variant (B.1.1.7) had replaced and became prevalent strains up to 46% in April of 2021. Unexpectedly, the Alpha variant was quickly reconstituted by the Delta variant (B.1.617.2) after half a year which become predominant and continued to spread globally till now. The Omicron (B.1.1.529) variant is spreading rapidly up to 70% isolates in Africa and 9% globally. Mutations on S protein raised one after another leading to mysteries whether such changes have a predictable regularity.

**Figure 1.**
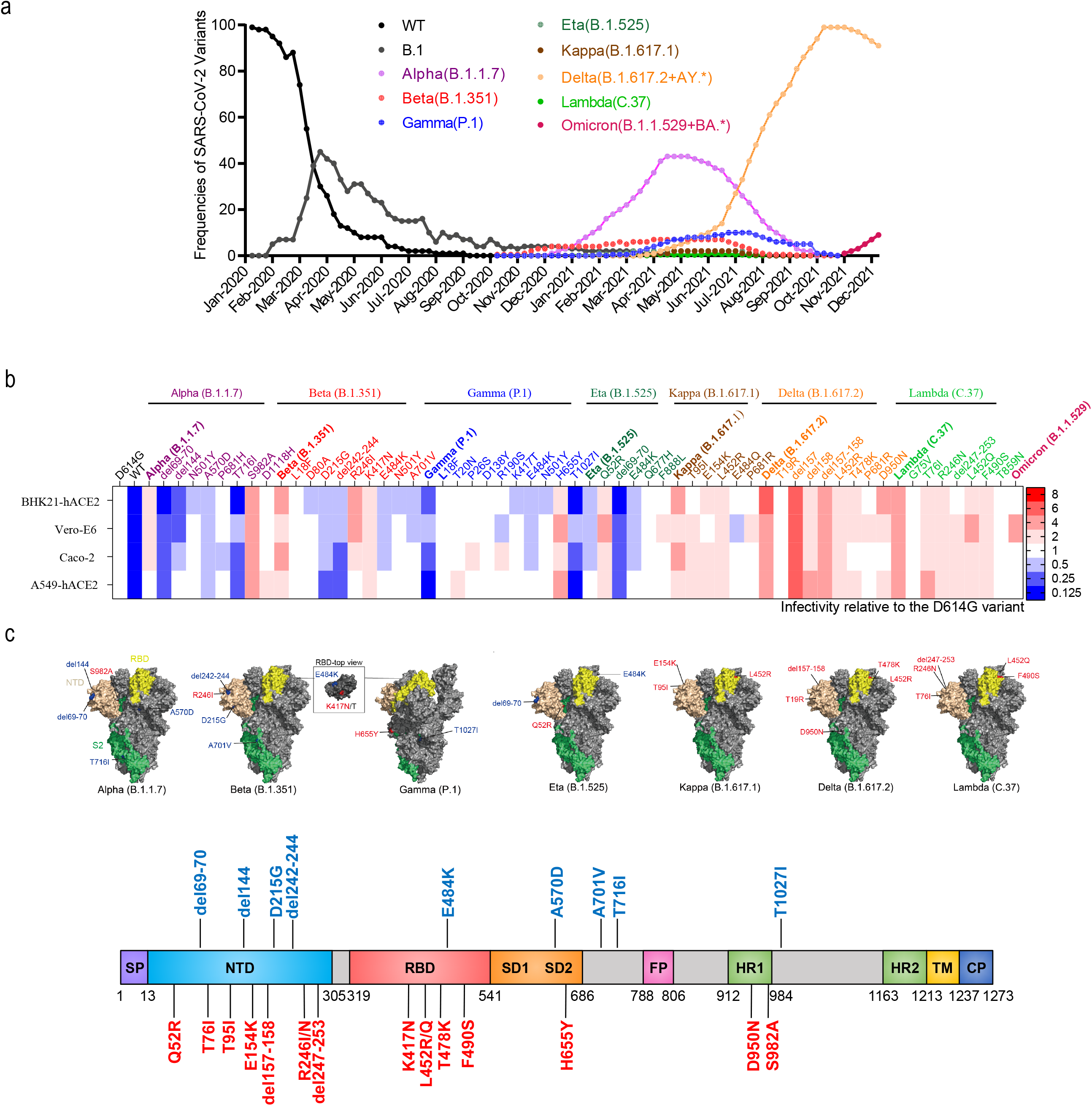
Infectivity analysis of SARS-CoV-2 variants. **a,** Line graph showing the frequencies of SARS-CoV-2 variants over time globally. The curve of cumulative variant counts was recorded according to the GISAID database. **b,** Infection assays with 54 SARS-CoV-2-related mutant pseudoviruses in four indicated cell lines, all of which are known to be susceptible to SARS-CoV-2. The infectivity of the D614G variant was set to 1. Blue and red represent decreased and increased viral infectivity to the reference D614G variant in the indicated cell lines, respectively. Data represent the means of three independent experiments. **c,** The location of crucial mutations affecting viral infectivity in spike protein. Crucial mutations were represented in both tertiary (PDB ID: 6VYB) and secondary structures of the spike protein. The color blue indicated decreased infectivity and red indicated increased infectivity.

To explore the functions of SARS-CoV-2 variants and mutants, we generated 54 pseudotyped viruses expressing either ancestry SWT or mutated S proteins (covering 45 single-site mutations and 8 defined variants generated in a 614G background) (illustrated in **Fig. S1**). The infectivity assays were performed with four SARS-CoV-2-susceptible cell lines: BHK21-hACE2, A549-hACE2, Vero E6, and Caco-2 (*26, 27*). As summarized in **Figure 1b** (also shown in **Fig. S2**), the Alpha (B.1.1.7), Beta (B.1.351), Kappa (B. 1.167.1), Delta (B.1.167.2), and Lambda (C.37) variants displayed 1.77-fold, 3.38-fold, 2.90-fold, 4.24-fold, and 4.09-fold increases in infectivity compared to the reference D614G variant, respectively (the D614G infectivity was set to 1). Of note, the Delta variant showed the highest infectivity not only with the combined mutations but also with the single-site mutation in this variant. By contrast, the Gamma variant (P. 1) and the Eta variant (B.1.525) exhibited 4.02-fold and 2.25-fold decreases in infectivity compared to the D614G variant, respectively. Remarkably, the emerging Omicron (B.1.1.529) variant displayed the highest increase (4.74-fold) in infectivity in the Vero E6 cell line but an average 1.58-fold decrease in the other three cell lines. As Vero E6 cell has been reported to support an endocytosis pathway for virus infection (*28*), the results indicate a probable preference for the endocytic pathway of the Omicron variant.

Referring to the individual single-site mutation, the del157 mutation in the Delta variant (B.1.617.2) and the S982A mutation in the Alpha variant (B.1.1.7) played epistatic effects to increase viral infectivity (5.70 and a 3.88-fold increase compared to the D614G variant, respectively). In addition, we found that several mutations synergistically increase infectivities within a certain variant, such as R246I and K417N in the Beta variant, E154K and L452R in the Kappa variant, and T76I, R246N, and L452Q mutations in the Lambda variant (**Fig. 1b**). On the contrary, some mutations counteracted each other, such as E484K and T1027I mutations neutralized the enhanced infectivity of H655Y to weaken viral infectivity of the Gamma variant (P.1) (Fig. 1b). Similarly, the E484K and del69-70 mutations also reduced Q52R infectivity in the Eta variant (**Fig. 1b**). In summary, single mutations in the NTD region from VOC or VOI could either increase or decrease the infectivity but mutations in the RBD region mainly enhance viral infectivity except for E484K (**Fig. 1c**). Interestingly, mutations in the S2 subunit, such as S982A and D950N in the HR1 region strongly enhanced viral infectivity indicating the S2 subunit also plays an essential role in regulating viral infectivity.

### SARS-CoV-2 variants exhibit decreased susceptibility to mAbs neutralization

To study the immune-escape abilities of S mutants, we test all the 54 pseudoviruses against a set of 14 monoclonal neutralizing antibodies targeting either WT S1 or RBD (**Table S1**, as a control, 14 of 27 commercial monoclonal antibodies could effectively neutralize the D614G variant in **Fig. S3**). Viral sensitivities to neutralization were calculated by neutralizing percentages relative to that of the reference D614G variant under the NT_50_-dilution to D614G variant for each antibody. Based on the relative neutralization activities, the 14 tested antibodies can be divided into four classes (**Fig. 2a left labeling**). Briefly, class I antibodies showed decreased neutralizing activity against almost all the single-site mutations and variants. Compared with class I, the antibodies in class II showed much higher neutralizing activities against several mutations in NTD and S2 such as L18F, del69-70, A701V, and T1027I (average neutralizing activities increased 1.28, 1.22, 1.61, and 2.08-fold compared with class I antibodies, respectively). However, class II antibodies showed little neutralization to the E484K mutation. On the contrary, the class III antibody exhibited a better response to the E484K mutation. Antibodies in class IV showed broad neutralization to a wide range of variants and single-site mutations.

**Figure 2.**
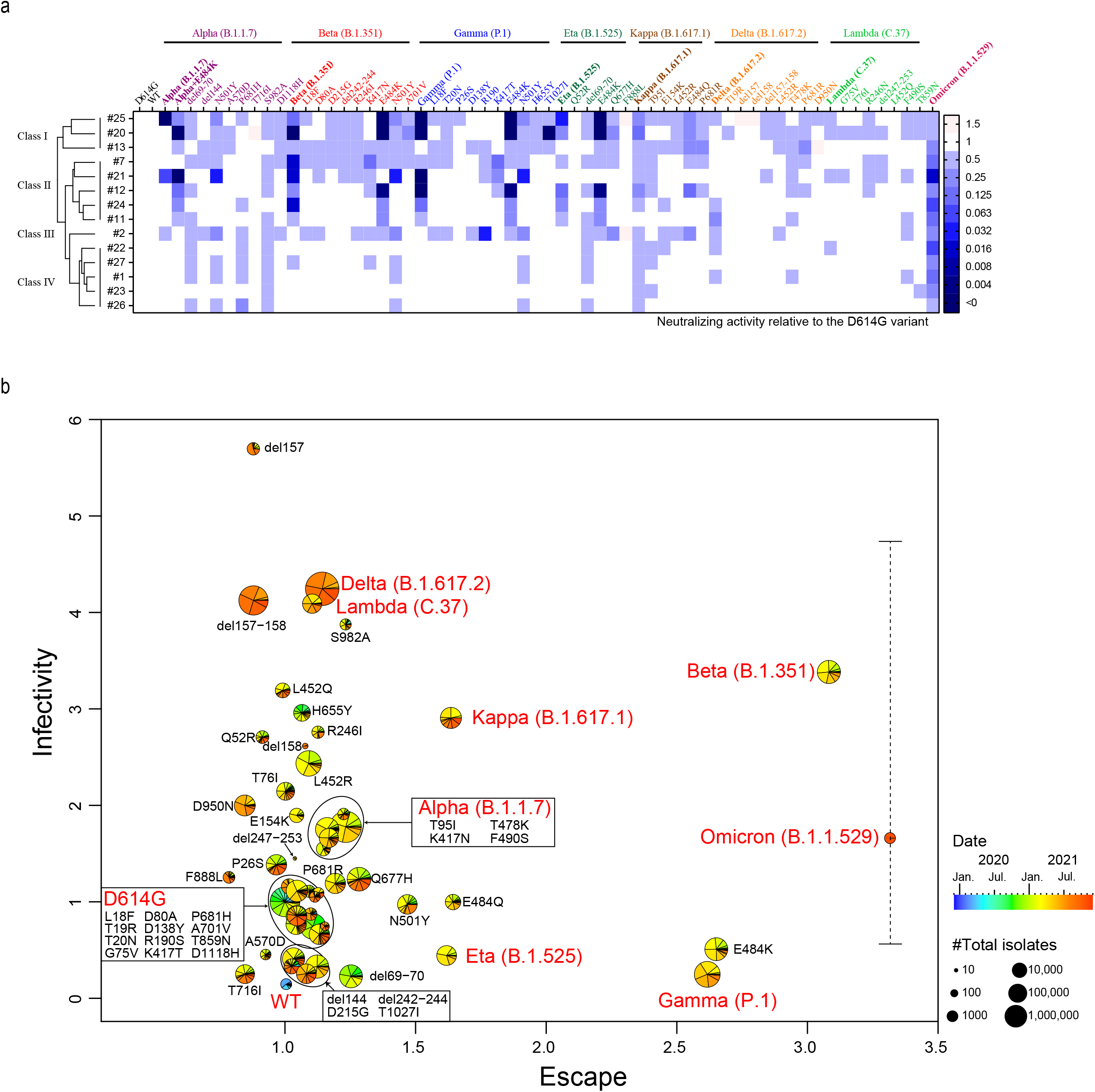
The neutralizing activity of the monoclonal antibody to SARS-CoV-2 variants and the evolutional tendency of SARS-CoV-2 spike protein. **a,** Heat map representation of neutralizing abilities using 14 neutralizing monoclonal antibodies against 54 SARS-CoV-2 related mutant pseudoviruses; the neutralization activity against the D614G variant was set as 1. Blue represents the decreased neutralization activity. Data represent the means of three independent experiments. **b,** The quadrantile diagram of SARS-CoV-2 evolution, X-axis show the immune-escape ability and Y-axis show the infectivity of SARS-CoV-2 variants (the error bar indicated the diversity of cell specific infectivity for Omicron). The color of pie charts indicated the isolation time of the indicated variants recorded in the GISAID database, the area size of each sector indicated numbers of isolates of individual variants until December 5^th^ 2021 (per month).

Examination on the single-site-mutant pseudoviruses showed that the most severe immune escape sites were E484K (2.65-fold reduction to the reference D614G variant in neutralizing activities against classes I-III antibodies) and N501Y (1.47-fold reduction to the reference D614G variant in neutralizing activities against classes I-III antibodies) (**Fig. 2a**). Of note, the E484Q in the Kappa variant showed a weaker immune-escape ability (~1.64-fold reduction to the reference D614G variant in neutralizing activities against classes I-III antibodies) as compared to E484K. This might be because the electric charge of E and Q resembles more than K. To prove E484K is crucial in immune-escape, we further introduced E484K mutation in the Alpha variant (B.1.1.7) (Alpha+E484K). The Alpha+E484K variant indeed showed a stronger immune escape than the Alpha variant (B.1.1.7) reaching up to ~30-fold more resistance in response to antibody #20. These results suggested that the amino acid on position 484, especially K484 in the RBD is crucial in viral immune-escape ability. Besides E484K/Q and N501Y, del69-70, S982A, and P681H also showed moderate immune-escape abilities, which should be considered in vaccine updating.

When referring to the variants, the average sensitivity of the Alpha variant (B.1.1.7) against monoclonal antibodies was similar to the D614G variant (**Fig. 2a**). The Beta variant (B.1.351) showed a high immune-escape level against classes I-III antibodies (3.08-fold reduction to the reference D614G variant in neutralizing activities). The Gamma (P.1) and Eta (B.1.525) variants also displayed a wider escape spectrum against classes I-III antibodies due to the E484K mutation (2.62 and 1.62-fold reduction respectively). Of concern, the Delta (B.1.617.2) and Lambda (C.37) variants only showed mild escape to all the four classes of antibodies. Strikingly, the emerging Omicron (B.1.1.529) variant showed the highest resistance among all the variants (3.32-fold reduction to the reference D614G variant in neutralizing activities against classes I-III antibodies) and even gained resistance to the class IV antibody (**Fig.2a, the far-right panel**).

### The evolutional tendency of SARS-CoV-2 spike protein

To understand the evolutional tendency of the SARS-CoV-2 spike protein, we generated a quadrantile diagram combined with isolates numbers to present the infectivity (Y-axis) and immune-escape (X-axis) of SARS-CoV-2 S protein based on the results in Figure 1b and Figure 2a. The color in each pie chart indicated the different isolation times of the individual variant (mutation), and the area size of each pie chart indicated numbers of isolates from January 2020 to December 5^th^ 2021 recorded in GISAID (the bigger size the larger number of isolates) (**Fig. 2b**). The comprehensive results from 4,710,001 S sequences showed that one set of mutations tends to increase the viral infectivity, such as del157, del157-158, S982A, H655Y, R246I, Q52R, and L452R, and another set of mutations tends to escape the antibodies such as E484K, N501Y, and E484Q. Interestingly, few mutations had a dual effect of affecting both infectivity and immune-escape ability of SARS-CoV-2.

Only the Beta variant distributed on the diagonal represented both high infectivity and high immune resistance. Of note, case numbers of the Beta variants were also small (**Fig. 2b**). Other variants with large numbers of isolates either evolved to have increasing infectivity (vertical) or high immune-escape ability (horizontal). For instance, the Delta and Lambda variants appear in the Y-axis direction with infectivity factors of 4.25 and 4.10, but they all showed low ability to escape neutralizing antibodies. On the contrary, the Eta and Gamma variants showed strong escape ability plus low infectivity appearing in the X-axis direction. The highly concerned Omicron variant is located close to the Beta variants with the highest immune-escape factor of 3.32, and high infectivity only in Vero E6 cells but low infectivity in the other three cell types. The data suggest that, in most cases, viral infectivity and immune escape ability of spike protein is contradictory. The highly infectious spike goes along with weak immune-escape ability, in contrast, the highly resistant variants work along with low infectivity. Although the prevalent situation for Omicron is still pending, the current evolutionary trend is similar to the Beta variant but it differed in cell-specific infectivity.

### Design a pan-vaccine with consensus S protein

Obviously from Figure 2b, the S protein evolved into two distinct evolutional paths. Thus, the current single-strain vaccine strategy could protect variants from the same evolutionary path but not from another path. For instance, the S_WT_ vaccine cannot protect the Beta or Gamma variants as efficiently as S_WT_ or the Alpha variant. To construct a pan-vaccine to elicit broad neutralizing antibodies to variants in both paths, we generated a consensus S protein by downloading 2674 Spike protein amino-acid sequences from the NCBI database (until February 28, 2021, before Delta emerged) to build a phylogenetic tree using MEGA 10.0 software (**Fig. 3a**). The S protein could be divided into five clades, of which clade 1 contains the most isolates of 2539 sequences, including variants from B.1 descendants and Gamma (P.1) variants. The Clade 2-3 contains 56 sequences, including a mixture of the Alpha (B.1.1.7), Beta (B.1.351), Eta (B.1.525) strains, etc. The Clade 4-5 contains 80 sequences, including mutants beyond the defined VOC or VOI nominates. Because the 2539 sequences in clade 1 are closely related and account for 95% weight of the database, we calculated one consensus sequence for clade 1, and then plus 136 sequences in clade 2-5 to obtain the frequency of concerning mutations in 137 sequences (**Figure 3b and Table S2**). Five mutations, D614G, del69-70, del144, N501Y, and P681H accounted for the top-five proportions (**Fig. 3b**). In addition, considering the results of the neutralization experiment in Figure 2a (E484K significantly resisted to neutralizing antibodies), we design a consensus sequence of S_6_ (E484K with above top-five-frequency mutations) (**Fig. 3c**) and another S_15_ (introducing 9 more top mutations based on S_6_) (**Fig. S4a, b**). We further reconstructed phylogenetic trees using the eight VOC variants (two isolate sequences each) and the S_6_ sequence in either a rectangle shape (**Fig. 3d**) or a clock shape (**Fig. 3e**). The results clearly showed that the S_6_ sequence is located in the middle of the phylogenetic tree indicating its consensus characteristic. Strikingly, although omicron emerged after we constructed the S_6_, it clustered together with the S_6_ sequence in the same clade despite a far evolutionary distance from all the other variants, proofing again that the S_6_ sequence is the root sequence taking the most consensus mutations (**Fig. 3d and 3e**).

**Figure 3.**
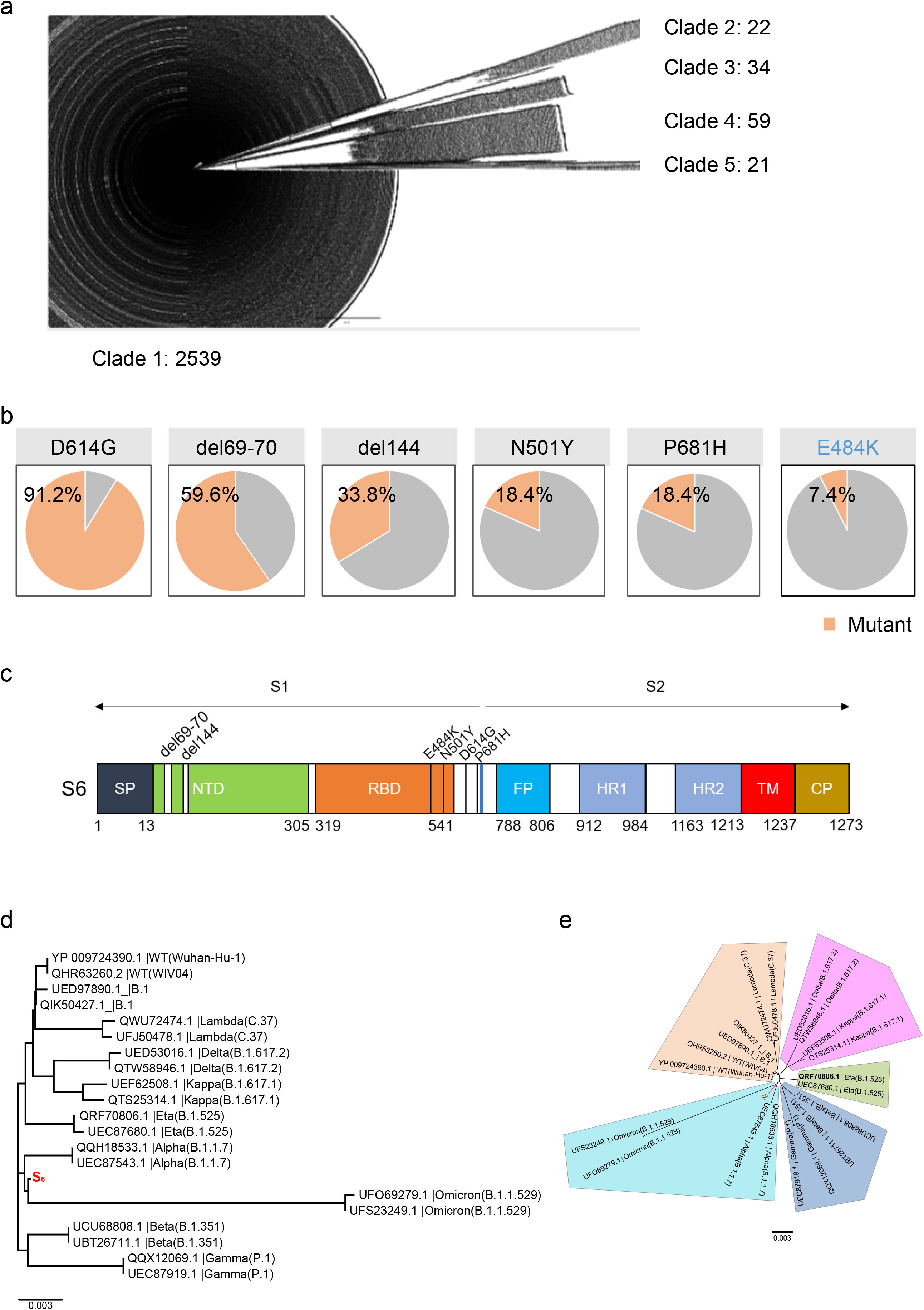
Design the consensus S_6_ sequence. **a.** Phylogenetic trees of 2674 spike protein amino acid sequences from the NCBI database. **b.** The proportion of top-five mutations and E484K mutations. **c.**Linear diagram of the full-length SARS-CoV-2 spike (S) updated protein showing the S_6_. **d and e.** The phylogenetic tree was constructed using the Neighbor-Joining method by MEGA 10.0 displayed either in a rectangle shape (d) or in a radiate shape (e). The S_6_ was highlighted in red (The line of S_6_ was magnified 10-fold for visualization).

During the time of preparing the manuscript. we had re-calculated the proportion of mutations base on the database update to Oct 2021 (when the Delta variant had already emerged) and ended up with the same top-five mutations in S_6_ (Data not shown).

### The universal S_6_ vaccine exhibits a broadly neutralizing ability to heterogeneous SARS-CoV-2 variants

To test whether S_6_ could serve as a pan-vaccine antigen against heterogeneous variants, we generated subunit vaccines by expressing S trimer of either S_WT_ or S_6_ in the CHO-K1 eukaryotic expression system. Subsequently, both S_6_ and S_wt_ antigens were mixed with adjuvants to formulate subunit vaccines, which were then used to immunize BALB/c or K18-hACE2 mice. Each mouse was immunized with 5 μg/dose antigen with primary and boost immunization strategy in an interval of 14 days (immunization procedure shown in **Fig. 4a**). The mouse sera were collected 14 days after the second vaccination.

**Figure 4.**
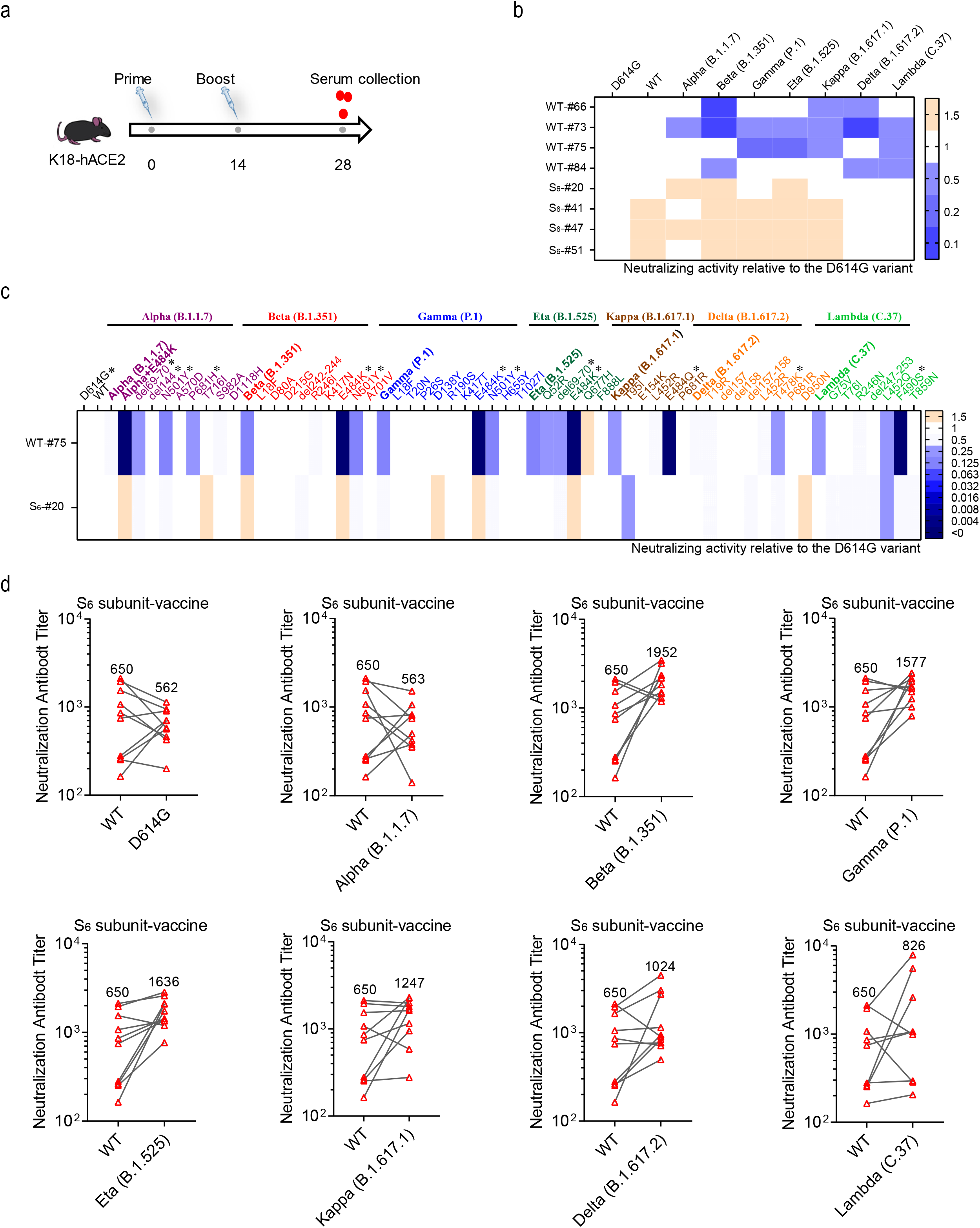
Neutralization of SARS-CoV-2 pseudoviruses by S_WT_ or S_6_ vaccinated mice serum. **a.** Schematic diagram of the vaccination protocol. The K18-hACE2 mice from each group were prime and boost-vaccinated either ancestry S_WT_ or S_6_ subunit vaccines (5 μg/dose) at day 0 and day 14. Sera were collected on day 28. **b.** Heat map representation of the neutralizing activities of ancestry S_wt_ (n=4) and S_6_ (n=4) vaccinated mice sera against VOC and VOI pseudoviruses. **c.**Heat map representation of neutralizing activities of one representative mice serum against all 54 pseudoviruses. For **b** and **c**, the neutralization activity against the D614G variant was set to 1. Blue and earthy yellow represent decreased and increased neutralizing activity of sera, respectively. Data represent the means of three independent experiments. The asterisk (*) indicated the mutations which showed more resistance to neutralization induced by ancestry S_WT_ as compared to S_6_. **d.** The geometric mean titer (GMT) analysis of the NT_50_ for S_6_ vaccinated mice sera against VOC and VOI pseudoviruses (n=10). The lines indicated the NT_50_ titer of the same sample with GMTs indicated by the labeling numbers. Data represent the means of three independent experiments.

To evaluate the immunogenicity of the S_wt_ and S_6_ vaccines, we first investigated the serum neutralizing activities from each vaccinated mouse to different pseudotyped variants under their NT_50_-dilutions to the reference D614G pseudovirus respectively. The results in **Figure 4b** showed that sera from the ancestry S_WT_ vaccinated mice sharply decreased their neutralizing activities to all the VOC or VOI variants, especially to the Beta variant (2.27-fold average reduction to the reference D614G variant) and Delta variant (1.94-fold average reduction to the reference D614G variant). However, sera from S_6_-vaccinated mice exhibited a broader spectrum of neutralizing activities to all the seven variants and showed even stronger neutralization to five out of seven variants (1.06- to 1.53-fold increased to the reference D614G variant) (**Fig. 4b**). A detailed serum neutralization against all the individual mutations in VOC and VOI showed that S_6_ sera could neutralize better against immune-escaping mutations such as E484K, N501Y, E484Q, del69-70, F490S, P681H, and T478K (**Fig. 4c**). We further evaluated NT_50_ of S_6_ vaccinated sera by calculating Geometric Mean Titers (GMT). The results in **Figure 4d** showed that S_6_-elicited NAbs neutralized WT pseudoviruses at an average GMT of 650. Meanwhile, it can also neutralize VOC and VOI pseudoviruses at the GMT range of 563-1952, indicating that the consensus S_6_ vaccine had universal protections to the antigen shift of SARS-CoV-2 variants.

To our surprise, the mice serum from the S_15_ vaccination did not show better neutralizing against different variants than the S_6_ (**Fig. S4**), which implies that the design of vaccine antigens is not a simple superposition of mutation sites but requires joint analysis of rigorous bioinformatics calculations and experimental evidence.

Taken together, the current ancestry S_WT_ vaccine with low neutralization to viral mutants needs to be updated to broad-reactive universal vaccines when facing the continuous antigen shit of SARS-CoV-2 variants.

### S_6_-vaccine protects K18-hACE2 mice from the WT, the Beta, and the Delta SARS-CoV-2 challenge

To assess the protective effect of S_6_ subunit-vaccine against authentic viruses, the K18-hACE2 mice (*29*) were immunized with 25 μg/dose S_6_ subunit-vaccine with a primary and boost immunization strategy. After vaccination, mice were grouped randomly (n=4-5) and challenged at lethal doses of either ancestral WT virus, or the Beta (B.1.351) virus, or the Delta (B.1.617.2) virus by intranasal infection (experimental procedure shown in **Fig. 5a**).

**Figure 5.**
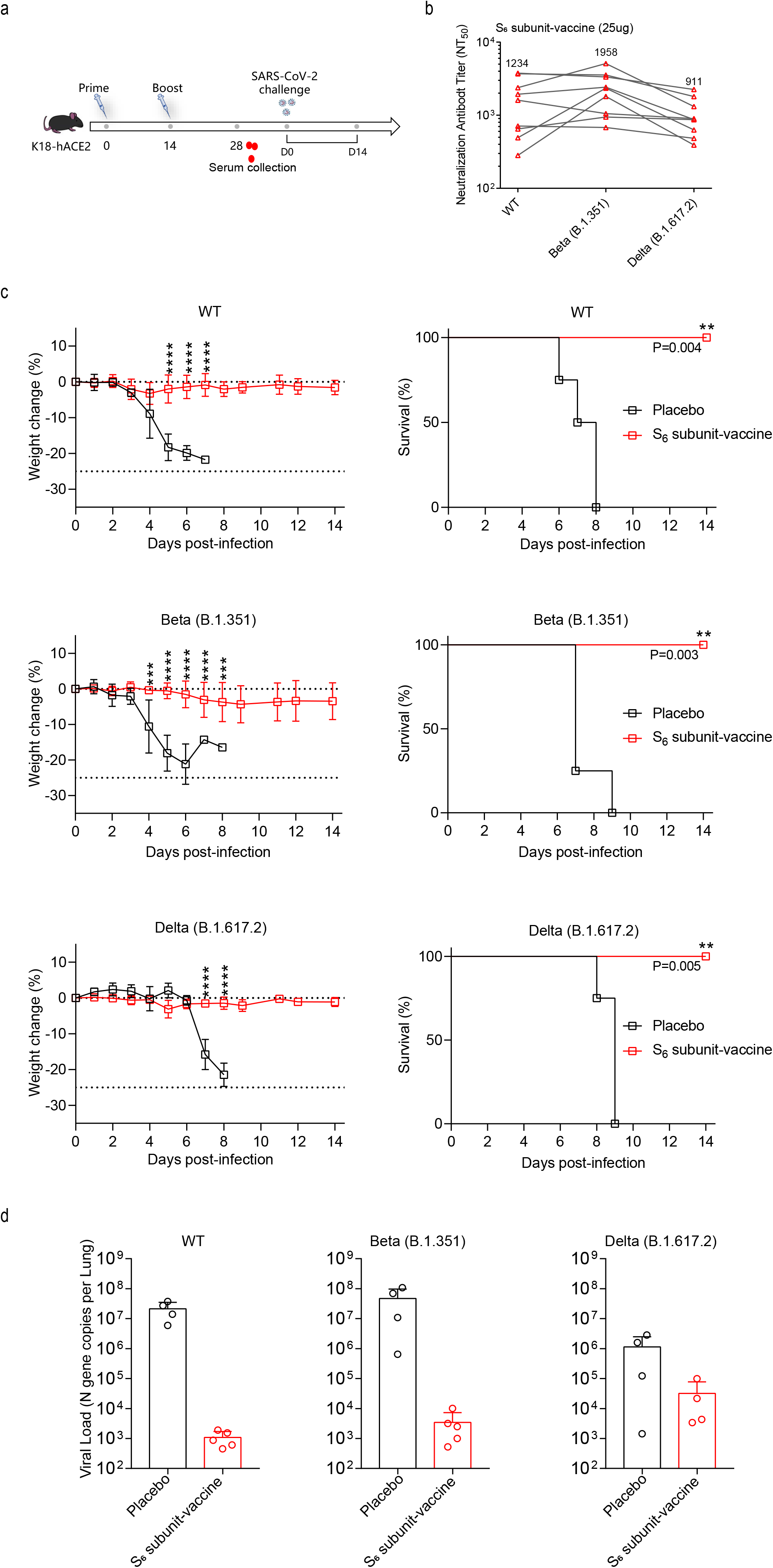
The S_6_ subunit-vaccine protects three SARS-CoV-2 variants infection in mice. **a.** Schematic diagram of the vaccination and virus challenging protocol. The K18-hACE2 mice from each group were prime and boost-vaccinated ancestry S_6_ subunit-vaccines (25 μg/dose) at day 0 and day 14. Sera were collected on day 28. After vaccination, mice were challenged with three different variants of SARS-CoV-2 by intranasal infection. The body weight and survival were monitored for 14 days or until body weights lost more than 25% (n=4-5 in each group). **b.** The geometric mean titer (GMT) analysis of the NT_50_ for S_6_ vaccinated mice sera against VOC and VOI pseudoviruses (n=10). The lines indicated the NT_50_ titer of the same sample. Moreover, the numbers indicated the GMTs. **c.** The body weighs and survival curves of mice described in a. **d.** The viral genome copy numbers of SARS-CoV-2 N were quantified. Values represent means ± SD of 4-5 individual mice. The body weights are present as the mean percentage of weight change ±SEM and survival curves are shown. *P < 0.05; **P < 0.01; and ***P < 0.001. Statistical analysis, two-way ANOVA for weight curves and bar graphs, and the log-rank test for survival curves.

The results in **Figure 5b** showed that the neutralization titers of mice sera against the pseudotyped WT, Beta, and Delta viruses were almost equally high at GMT of 1234, 1958, and 911, respectively (**Fig.5b**). The body weights of mice from the unvaccinated control group all dropped to more than 25% and died on day 6 to day 9. In contrast, the S_6_ subunit-vaccine could provide 100% protection against all the three viruses (WT, Beta, and Delta) (**Fig.5c**). Consistent with the body weight and survival data, S_6_ subunit-vaccine reduced viral RNA load at 19677, 13650, and 36-fold in the lung of the WT, Beta, and Delta SARS-CoV-2 infected mice compared to the control group respectively (**Fig.5d**).

Overall, our results confirmed that S_6_ subunit-vaccine conferred broad protection against SARS-CoV-2 variants evolved from two distinct paths. It can also elicit equal effective NAbs against the emerging Omicron variant at the GMT of 1214 (**Fig. 6**), suggesting that the consensus S_6_ immunogen can most likely serve as a pan-vaccine to protect against potential antigen drift of SARS-CoV-2 in the future.

**Figure 6.**
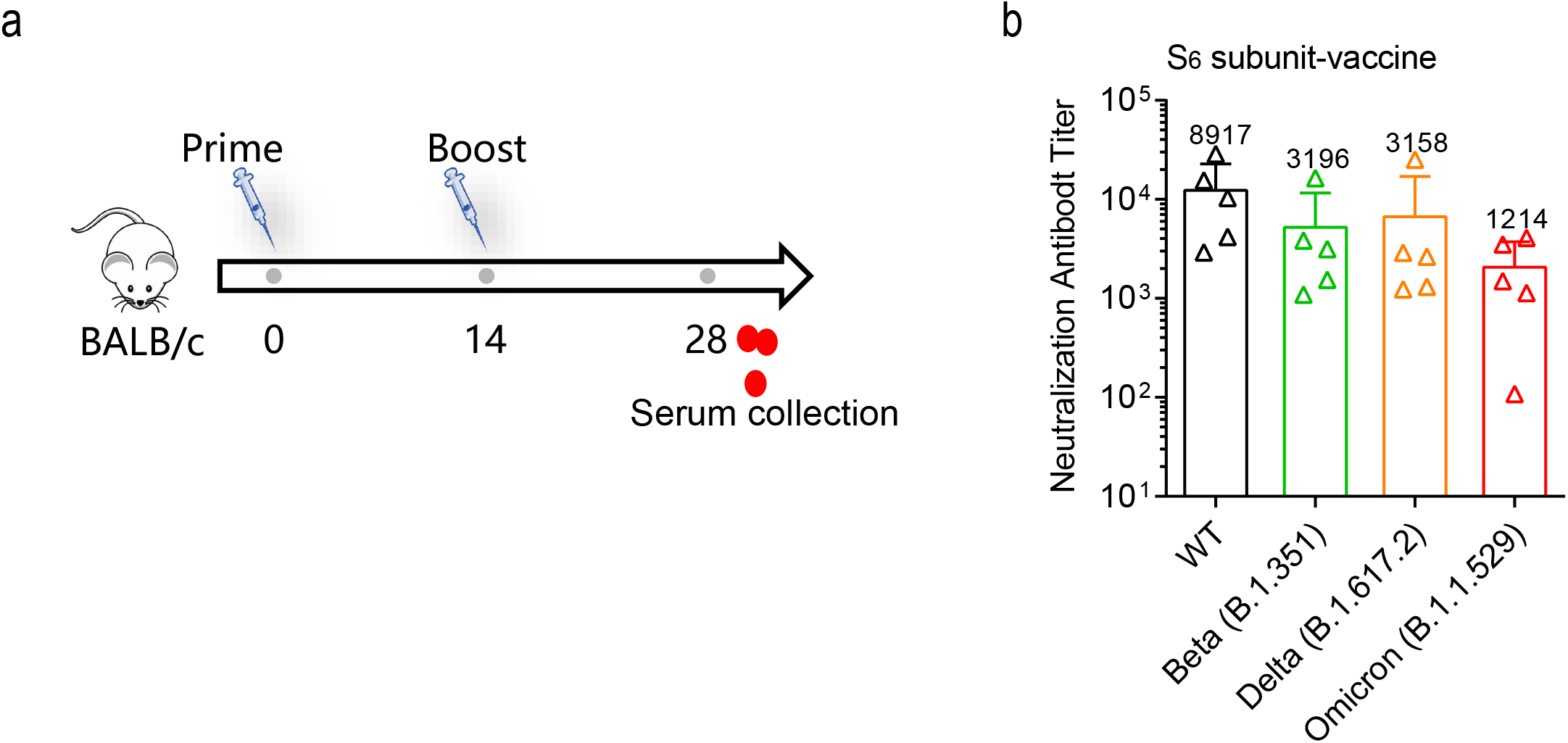
Immunogenicity of S_6_ subunit-vaccine against the Omicron (B.1.1.529) variant. **a.** Schematic diagram of the vaccination. The BALB/c mice were prime and boost-vaccinated S_6_ subunit vaccines (25 μg/dose) at day 0 and day 14. Sera were collected on day 28. **b.** The geometric mean titer (GMT) analysis of the NT_50_ for S_6_ vaccinated mice sera against the WT, Beta, Delta, and the Omicron (B.1.1.529) pseudoviruses (n=5 for each group). The numbers indicated the GMTs. Data represent the means of three independent experiments.

## Discussion

The emergence of a variety of SARS-CoV-2 variants has become a major concern during the pandemic. More than 6,200 types of amino acid substitutions, deletions, and insertions have been found in the SARS-CoV-2 S protein. Mutations are categorized into two directions. One makes the virus more infectious with high cell binding and entry ability which leads to more transmissible in the human population. The other enables the virus to evade the host’s immune responses such as neutralizing antibodies leading to breakthrough infections after vaccination (*26, 30, 31*).In particular, L452R has been reported to increase the infectivity of the B.1.617 variant (*32*). For immune-escape ability, E484 is the most important amino acid. Changes of E484 to K, Q, or P reduced the neutralizing activity of antibodies to more than 10 folds (*33*). How SARS-CoV-2 evolves in the future is still mysteries and is far from certain.

In this study, the infectivities of 54 pseudotyped SARS-CoV-2 viruses taking individual mutations were systemically studied. We found that the significantly effective mutations mainly enriched in NTD and RBD, while mutations outside of these two domains had minimal effects on viral infectivity (Fig 1c). To our surprise, S982A mutation, which is located in the heptad repeat 1 (HR1) domain of the S2 subunit, showed a high ability to increase viral infectivity. The HR1 domain was highly conserved among HCoVs and play an essential role during the viral membrane fusion process, serving as a broad-spectrum drug target (*34, 35*). Besides, S982A exerts decrease sensitivities to neutralizations against Class IV antibodies which are broad reactive to other mutations, indicating that S982A mutation should be considered in both enhanced transmissibility and drug resistance.

The results from our immune escape analysis are similar to published studies that the Beta (B.1.351) and Gamma (P.1) variants showed strong resistance against neutralizing antibodies. Particularly, both the Beta (B.1.351) and Gamma (P.1) variants had E484K and N501Y mutations in the RBD that contributed to evade immunity. Interestingly, E484K and N501Y cannot increase viral infectivity to the reference D614G mutant, suggesting that infectivity and immune escape ability of a certain mutation are incompatible. Nevertheless, combinations of several mutations in the Beta variant increased both the infectivity (contributed by R246I and K417N) and immune escape ability (contributed by E484K and N501Y) to make more significant gains. However, such variants gaining dual functions as the Beta variants are limited with a relatively small number of isolates. On the contrary, the Delta variant with a large isolate number showed the highest infectivity but only a mild immune-escape ability, indicating that circulating SARS-CoV-2 strains might obtain higher infectivity by sacrificing a strong immune-escape ability. Unexpectedly, the Omicron variant emerged during our manuscript preparation. The analysis of Omicron showed the strongest immune escape even than the Beta variant probably contributed by the most mutations it takes. However, the infectivity of Omicron is not the same as the Beta variant which exhibited an average increase in all the four tested cell lines. The Omicron variant was significantly more infectious in Vero E6 cells but less infectious in BHK21-hACE2, A549-hACE2, and Caco-2 cells. We suspect that this emerging variant Omicron might still struggle in a balance between infectious and evading immunity so that the evolutionary path of the Omicron variant is still uncertain either like the Beta or the Gamma variant.

SARS-CoV-2 mainly depended on two pathways (TMPRSS2 mediated membrane fusion pathway and cathepsin B/L mediated endocytic pathway) to enter cells (*36*). Previous studies showed that SARS-CoV-2 entry into the Vero E6 cells mainly through the endocytic pathway due to the low expression of TMPRSS2 in this cell. This might indicate that the Omicron variant prefers the endocytic pathway to enter cells. It has shown that TMPRSS2 is highly expressed in alveolar type I and II cells of the lung (*37*) so that the Omicron variant might be relatively less infectious in these cells. More studies are demanded on the transformation of susceptible cell types because of the changes in the viral entry pathway.

As the SARS-CoV-2 spike protein evolved in two distinct directions (high infectivity with weak immune-escape or high immune-escape with low infectivity) (Fig 2b), a universal vaccine to provide broad protection must take all the heterogeneous mutations into account within the same antigen. Here, we generated a consensus antigen (S_6_) by phylogenetically calculating the consensus sequence which is a collective sequence including the most frequent mutations. The S_6_ antigen included the top five high-frequency mutations and the most resistant mutation (E484K). The high-frequency mutations in the S_6_ antigen protected against variants with high infectivities like the Delta and Lambda variants, while the protection of high immune-escape variants was correlated with the E484K mutation. Our results showed that the S_6_ antigen could elicit broad humoral immune against circulating variants including the WT, the Beta (B.1.351), and the Delta (B. 1.617.2) variants. Interestingly, our S_6_ antigen not only improved the protective effects of covered mutations but also elicited better neutralization activity against uncovered mutations like (E484Q, T478K, and F490S) (Fig 4c). Consistent with this, class III and IV antibodies also showed broad-spectrum neutralization activity, suggesting that conserved epitopes might exist in SARS-CoV-2 spike protein. We further test the protection of the S_6_ vaccine against the Omicron variant. Our results showed that three out of four sera of the S_6_-vaccinated mice displayed similar neutralization against the Omicron variant as compared to the WT strain, and only one S_6_-serum decreased its activity. Recent studies have shown that the neutralizing activity of sera from ancestral strain vaccine generally decreased around 22 to 41-fold against the Omicron variant (*38, 39*), but the majority of our S_6_ sera performed better neutralization once again proofing the consensus characteristics of the S_6_ vaccine.

There was a rapid development of various types of vaccine such as mRNA vaccine (*40*), viral vector-based vaccine (*41*), inactivated vaccine (*42*), virus-like particle vaccine (*43*), and subunit vaccine (*44*). However, the current single-strain vaccine strategy with all these vaccine types could not provide cross-variant neutralizations especially to variants from two evolutionary paths. According to the current reports, all the ancestral S_WT_ vaccines decrease their neutralizations to the highly evading Beta and Gamma variants (*45, 46*). Convincingly, our consensus pan-vaccine strategy paves a way to fight against antigen drift of SARS-CoV-2 with great potentials.

## Materials and methods

### Cell lines

The A549-hACE2, Caco-2, Vero E6 cell lines were obtained from ATCC and maintained in Dulbecco’s modified Eagle’s medium (DMEM; Gibco) supplemented with 10% fetal bovine serum (FBS). BHK21-hACE2 cell line was provided by Professor Huan Yan, Wuhan University. All cells were incubated at 37 °C in 5% CO_2_.

### Viruses and mouse infection

The WT, Beta, and Delta SARS-CoV-2 viruses were kindly supplied by Hubei CDC and multiplied in Vero E6 cells to make virus stock. For mouse infection, 250PFU of WT, Beta, and Delta viruses were intranasally infected K18-ACE2 mice respectively in 50ul PBS in the A3 laboratory of Wuhan University, and the mice were then monitored daily for their body weights and survivals. The corresponding animal experiments have been approved by the Animal Committee of Wuhan University.

### Plasmid and Site-directed mutagenesis

The DNA sequences of human codon-optimized S proteins from SARS-COV-2 variants (B.1.1.7, GISAID: EPI-ISL-601443; B.1.351, GISAID: EPI_ISL_678597; P.1, GISAID: EPI_ISL_906075; B. 1.525, GISAID: EPI_ISL_1093472; B.1.2, GISAID: EPI_ISL_2000505; B. 1.617.1 GISAID: EPI_ISL_1660428 S1; B.1.627.2, GISAID: EPI_ISL_2029113; C.37, GISAID: EPI_ISL_3023383; B.1.1.529, GISAID:EPI_ISL_7162071) and S protein mutations were commercially synthesized or generated by overlapping-PCR based mutagenesis using pCAGGS-SARS-CoV-2-S-C9 (gifted from Dr. Wenhui Li, National Institute of Biological Science, Beijing, China) as a template and cloned into pCAGGS vector with C-terminal 18 aa truncation to improve VSV pseudotyping efficiency (*47*).

### Pseudovirus production

Pseudotyped VSV-ΔG viruses expressing either a luciferase reporter or mCherry reporter were provided by Professor Ningshao Xia, Xiamen University. To produce pseudotyped VSV-ΔG-Luc bearing SARS-CoV-2 spike protein (pseudo-SARS-CoV-2), Vero E6 cells were seeded in 10-cm dishes and transfected simultaneously with 15 μg SARS-CoV-2-S-Δ18 plasmid using Lipofectamine 2000 (Thermo). Forty-eight hours post-transfection, 500 μl pseudotyped VSV-ΔG bearing VSV-G protein was used to infect Vero E6 cells. Cell supernatants were collected after another 24 h, clearing of cell debris by centrifugation at 3000 rpm for 6 minutes, aliquoted, and stored at −80 °C.

### Titration of the pseudoviruses

The day before pseudo viruses titration, BHK21-hACE2 cells were seeded into 96-well culture plates at an appropriate density. To titrate the pseudoviruses, an initial 10-fold dilution in infection medium (DMEM, supplemented with 2% FBS, 1% penicillin/streptomycin) with six replicates were made in 96-well plates followed by serial 3-fold dilutions for a total of nine dilutions. The last column of the 96-well plate was served as the cell control without pseudoviruses. Then, the diluted pseudoviruses were transferred to the target cells and incubated at 37 °C in a humidified atmosphere with 5% CO_2_. After incubation for 24 h, the supernatant was sucked out and the cells were washed with cold PBS, followed by lysing in 40 μL cell lysis buffer (Promega, E1941). Then, 20 μL of the lysate and 10 μL of the luciferase substrate (Promega, E4550) were added into white 96-well plates for chemiluminescence detection using a Varioskan LUX Multimode Microplate Reader (ThermoFisher, VLBLATGD2). The 50% tissue culture infectious dose (TCID_50_) of the pseudoviruses is calculated according to the Reed-Muench method. In addition, wells with a relative luminescence unit (RLU) value equal to or higher than 10000 were considered positive (*48*).

### Pseudovirus infection assay

The day before pseudoviruses infection, the targeted cells were seeded into 96-well culture plates at an appropriate density. Using the lowest TCID_50_ as a reference, the pseudoviruses were diluted to the same TCID_50_ with the infection medium. After normalization, 100 μL of the pseudovirus with 2-fold dilution was added into the targeted wells in 96-well cell culture plates. After 24 h incubation at 37 °C with 5% CO_2_, the cells were washed with cold PBS and lysed in 40 μL cell lysis buffer. The chemiluminescence detection, as described in the titration of pseudoviruses, was performed to detect the RLU values. Each group contained three replicates and these experiments were performed three times independently.

### Neutralization assays

The day before neutralization assays, the BHK21-hACE2 cells were seeded into 96-well culture plates at an appropriate density. The SARS-CoV-2 pseudoviruses were incubated with serial-diluted sera at 37□ for 1h in 96-well white flat-bottom culture plates and then mixed with BHK21-hACE2 cells. After 24h, the cells were lysed in 40 μL cell lysis buffer (Promega, E1941). Then, 20 μL of the lysate and 10 μL of the luciferase substrate (Promega, E4550) were added into white 96-well plates for chemiluminescence detection using a Varioskan LUX Multimode Microplate Reader (ThermoFisher, VLBLATGD2). The 50 % neutralization dilution titer (NT_50_) and Geometric Mean Titers (GMT) with NT_50_ were calculated by GraphPad Prism 7 (GraphPad Software, Inc., San Diego, CA). Software with nonlinear regression curve fitting (normalized response, variable slope).

### SARS-CoV-2 S protein expression and purification

The DNA sequence encoding SARS-CoV-2 S residues 1–1208 (GenBank MN908947 nucleotides 21563-25384) was expressed in the Chinese hamster ovary (CHO-K1) cells by codon optimization. We constructed the sequences carrying 6 mutations (S_6_) and 15 mutations (S_15_) through rigorous bioinformatics calculation. The sequences were synthesized and constructed between Hind III and Not I site into GS-KS001 plasmid (Quacell biotechnology). Then, the CHO-K1 cells were transfected with the S_wt_, S_6_, or S_15_ plasmids by Bio-Rad Gene Pulser Xcell system, respectively. The S_wt_ protein was purified by Ni-NTA columns (Genscript) The S_6_ or S_15_ proteins were purified by anion-exchange column (Tosoh) and Hitrap Capto Core400 (Thomas Scientific). The purified S_wt_, S_6_, or S_15_ proteins were quantified by ELISA assay (Thermo Fisher Scientific) and stored at −80□.

### Immunization of mice

Female K18-hACE2 transgenic mice were obtained from GemPharmatech Co., Ltd, Jiangsu Province. For antigen formulation, 5 μg S_wt_, S_6_, or S_15_ proteins were mixed with an equal volume of Al(OH)_3_ for each mouse, respectively. The K18-hACE2 or BALB/C mice aged 6–7 weeks were immunized twice by intramuscular injection (100 μL in each of the left and right quadriceps femoris muscles per mouse) at 2 weeks apart. Total serum anti-S IgG titer was detected by ELISA assay using custom 96-well plates coated with S protein purchased from SinoBiologic.

### Crystallization and structure determination

The SARS-CoV-2 S-trimer profile (PDB: 6XR8) was downloaded from Protein Data Bank (PDB). The structural figures were obtained using the PyMOL Molecular Graphics System, Version 1.0.

### Evolutionary dynamics of SARS-CoV-2 variants

Evolutionary dynamics of SARS-CoV-2 variants from the ongoing COVID-19 pandemic were available at https://www.gisaid.org/phylodynamics/global/nextstrain/. In brief, the visualization only contained 3815 genomes in a single view for performance and legibility reasons (*49, 50*). The frequencies of SARS-CoV-2 variants in the first week of each month were recorded and the analysis was performed using Graph Pad 8.0.

### Defined the emerging time of single mutations

The peptide sequences of SARS-CoV-2 spike protein were downloaded from GISAID till December 5^th^ 2021. The original data exactly matched the “FASTA header format” and was used in the further analysis. MAFFT (v7.487) was used to align each peptide sequence to the reference sequence (Wuhan-Hu-1/2019), and the mutation information was parsed using a custom python script. We analyzed 46 kinds of monomer mutations contained in 7 kinds of combined mutants and classified each sequence according to the submission date which indicated the occurrence of these monomeric mutations (ignored mutations in other locations). To improve the accuracy, a sequence was discarded if it misses sequence information in any of annotated monomer mutation sites. For the statistics of the appearance time of the wild-type sequence, we required it to be completely consistent with the reference sequence.

## Acknowledgments

This work was supported in part by the National Key R&D Program (2018FYA0900801 and 2018FYA0900803 to K.X.), the Innovation Team Research Program of Hubei Province (2020CFA015 to K.L. and K.X.), the National Natural Science Foundation of China (grants 31922004 and 81772202 to K.X.), Application & Frontier Research Program of the Wuhan Government (2019020701011463 to K.X.). We are grateful to a special fund for COVID-19 Research of Wuhan University to K.L., the fund from Taikang Insurance Group Co., Ltd and Beijing Taikang Yicai Foundation, and the Fundamental Research Funds for the Central Universities for their great supports of this work.

We thank for technological supports from members in the SARS-CoV-2 Vaccine Task Force Group for their efforts to support the vaccine development, they are Dixiao Tang, Liang Qu, Yunxia Xu, Qianyun Liu, Xianying Chen, Ming Guo, Xin Wang, Zhixiang Huang, Ming Dai, Xiaoying Wu, Qing Xiong, Yifan Zhong, Ruichen Sun, Yingjian Li, Fang Liu, Chanjuan Zhou, and Yan Zhang.

## Author Contributions

K.L. and K.X. conceived the project and designed the experiments. YL.Z., L.D., W.N., S.L., W.Z., D.Z., D.N., M.X., Q.Z. constructed the mutation plasmids of SARS-CoV-2 Spike protein and produced the pseudoviruses. L.D., D.N., Q.Z. titrated and evaluated the infectivity of pseudoviruses. K.X., S.L., Z.C. designed the pan-vaccine and S.L. calculated the consensus sequence. W.N., Z.C. performed vaccine immunization of mice and collected the sera. YL.Z., W. N., M.X., Z.C., W.G. elucidated the immune escape of pseudoviruses against monoclonal antibodies and vaccine sera. Y.Z., D.W. performed the bioinformatics analysis. H.Y., Y.C., Y.P., X.W., Y.Y., J.L., Y. J. provided technical support and materials. YL.Z., L.D., Z.C., K.X., K.L. wrote the manuscript with input from all the other authors. YC.Z., S.L., Z.Z. performed the live virus experiments in the ABSL-3 lab. All authors read and approved the final manuscript.

## Notes

### Competing Interest Statement

The authors have declared no competing interest.

