## supplementary materials for "Construction of a potent pan-vaccine based on the evolutionary tendency of SARS-CoV-2 spike protein"

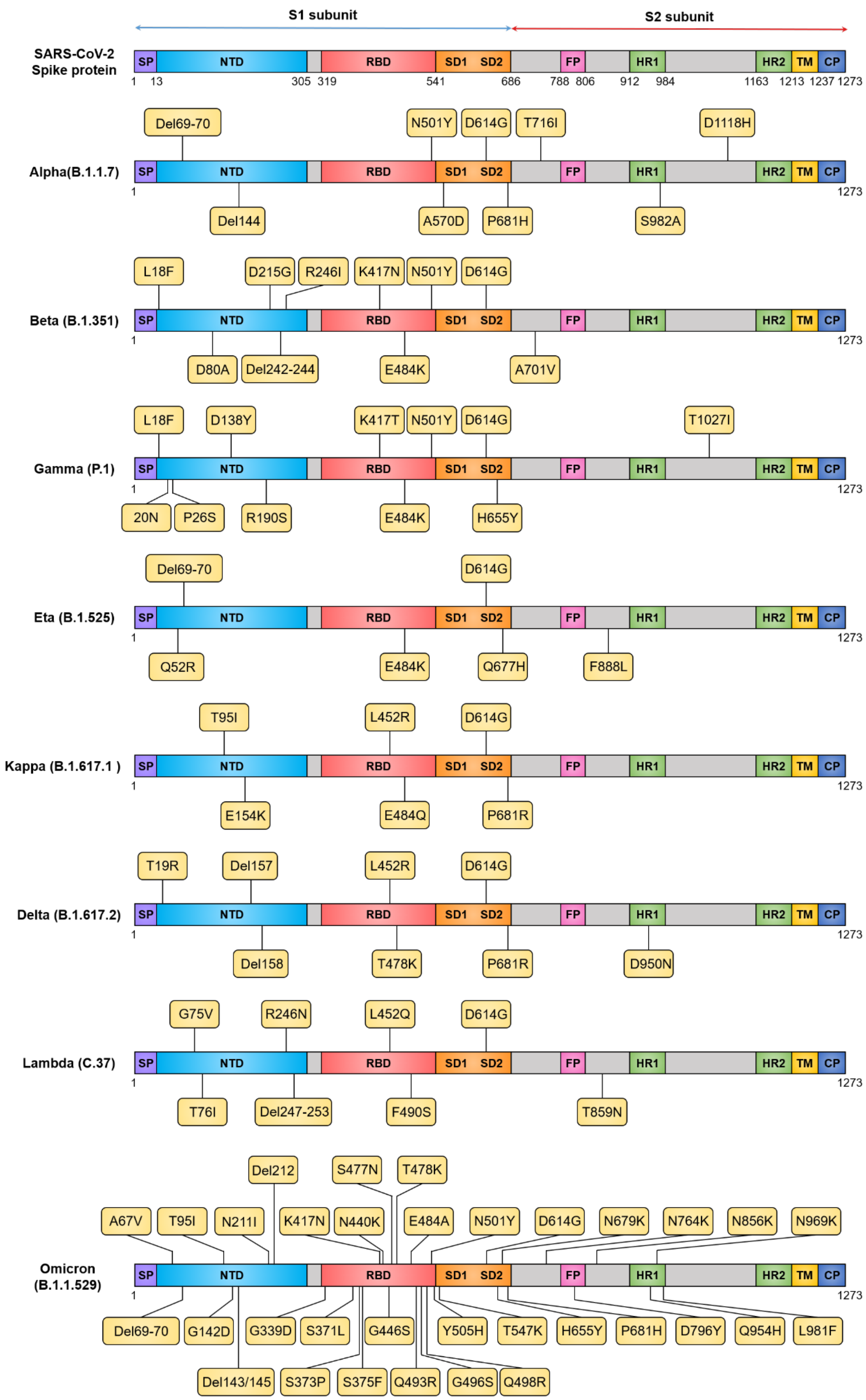

S1 subunit
S2 subunit
SARS-CoV-2 Spike protein
SP
NTD
RBD
SD1
SD2
FP
HR1
HR2
TM
CP
1 13 305 319 541 686 788 806 912 984 1163 1213 1237 1273
Alpha(B.1.1.7)
Del69-70
N501Y
D614G
T716I
D1118H
Del144
A570D
P681H
S982A
Beta (B.1.351)
L18F
D215G
R246I
K417N
N501Y
D614G
D80A
Del242-244
E484K
A701V
Gamma (P.1)
L18F
D138Y
K417T
N501Y
D614G
T1027I
20N
P26S
R190S
E484K
H655Y
Eta (B.1.525)
Del69-70
D614G
Q52R
E484K
Q677H
F888L
Kappa (B.1.617.1 )
T95I
L452R
D614G
E154K
E484Q
P681R
Delta (B.1.617.2)
T19R
Del157
L452R
D614G
Del158
T478K
P681R
D950N
Lambda (C.37)
G75V
R246N
L452Q
D614G
T76I
Del247-253
F490S
T859N
Omicron (B.1.1.529)
Del212
S477N
T478K
A67V
T95I
N211I
K417N
N440K
E484A
N501Y
D614G
N679K
N764K
N856K
N969K
Del69-70
G142D
G339D
S371L
G446S
Y505H
T547K
H655Y
P681H
D796Y
Q954H
L981F
Del143/145
S373P
S375F
Q493R
G496S
Q498R

**Figure S1.** Illustration of mutant residues along the S protein identified in SARS-CoV-2 variants. Mutant residues in each variant are represented by a black rectangle with a yellow background.

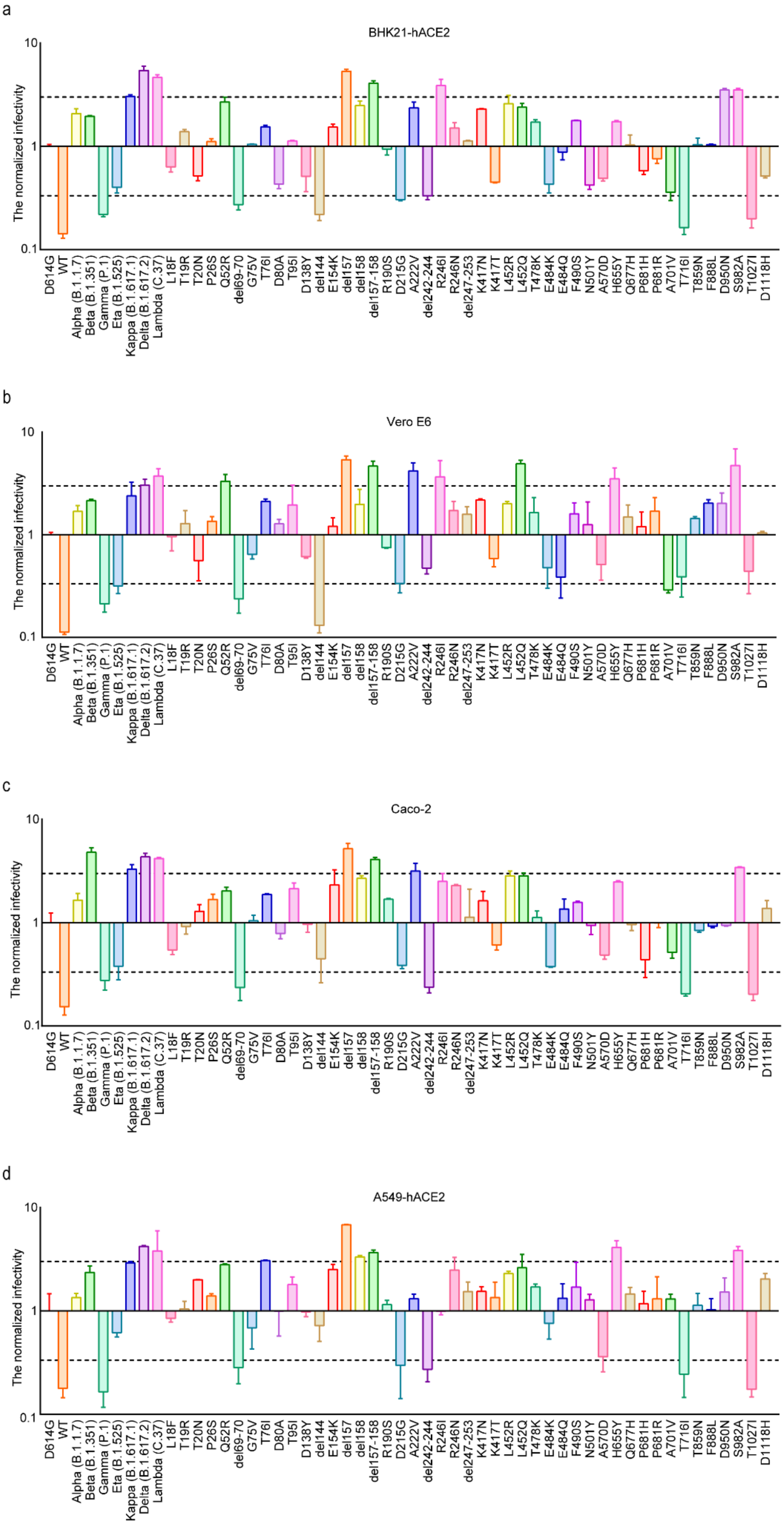

a
BHK21-hACE2
b
Vero E6
c
Caco-2
d
A549-hACE2
The normalized infectivity
10
1
0.1
D614G
WT
Alpha (B.1.1.7)
Beta (B.1.351)
Gamma (P.1)
Eta (B.1.525)
Kappa (B.1.617.1)
Delta (B.1.617.2)
Lambda (C.37)
L18F
T19R
T20N
P26S
Q52R
del69-70
G75V
T76I
D80A
T95I
D138Y
del144
E154K
del157
del158
del157-158
R190S
D215G
A222V
del242-244
R246I
R246N
del247-253
K417N
K417T
L452R
L452Q
T478K
E484K
E484Q
F490S
N501Y
A570D
H655Y
Q677H
P681H
P681R
A701V
T716I
T859N
F888L
D950N
S982A
T1027I
D1118H

**Figure S2.** Infectivity analysis of SARS-CoV-2 mutant pseudoviruses. **a-d**. Infection assays with the 7 crucial SARS-CoV-2 variants and 45 single-site mutants pseudoviruses with the four indicated cell lines, all of which are known to be susceptible to SARS-CoV-2.

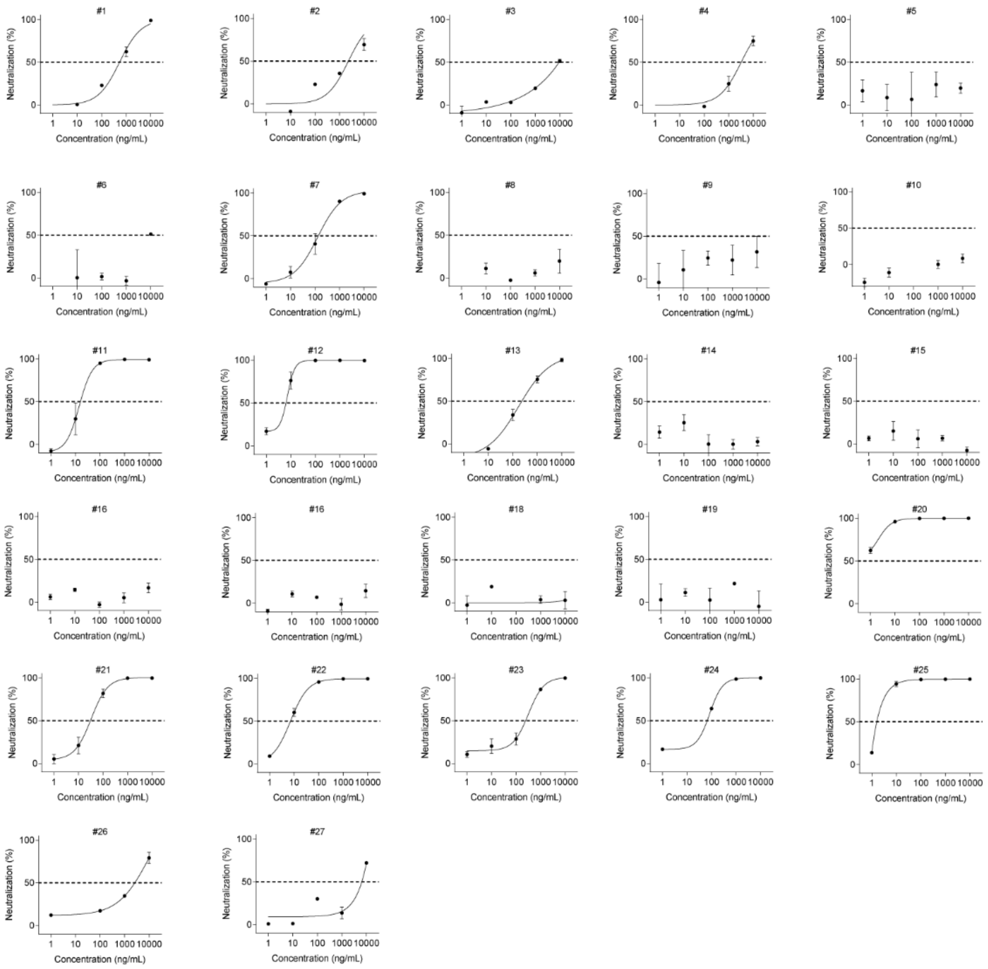

**Figure S3.** Neutralization of 27 monoclonal antibodies against the D614G variant pseudovirus, related to **Figure 2a**.

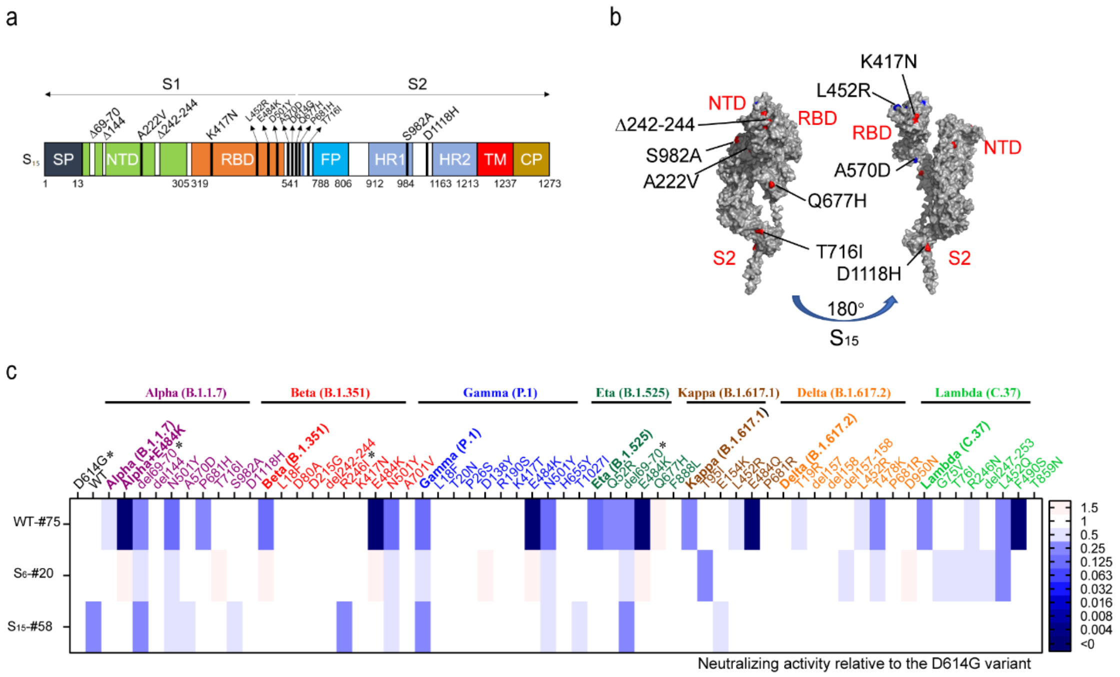


**Figure S4.** Neutralization of SARS-CoV-2 variant pseudoviruses in mice serum samples, related to **Figure 3e**

**a.** Linear diagram of the full-length SARS-CoV-2 spike (S) updated protein showing the $S_{15}$. **b.** Structure-based mapping of SARS-COV-2 spike (S) updated variants on the wild-type spike protein trimer (PDB ID: 6XR8). The variants are distributed on the surface of the complex. **c.** Heat map representation of neutralization reactions using the sera of mice that were injected with ancestry, $S_6$ or $S_{15}$ recombinant protein vaccine(n=1/group); the ratio of inhibition value (for each of the tested mice sera) detected for the reference D614G variant to the inhibition value for each of SARS-CoV-2-related mutant pseudoviruses. Blue and pink represent decreased and increased viral sensitivity to mice serum neutralization, respectively. The asterisk (*) indicated the mutations which showed more neutralization resistance compared to $S_{15}$ with $S_6$. Data represent the means of three independent experiments.

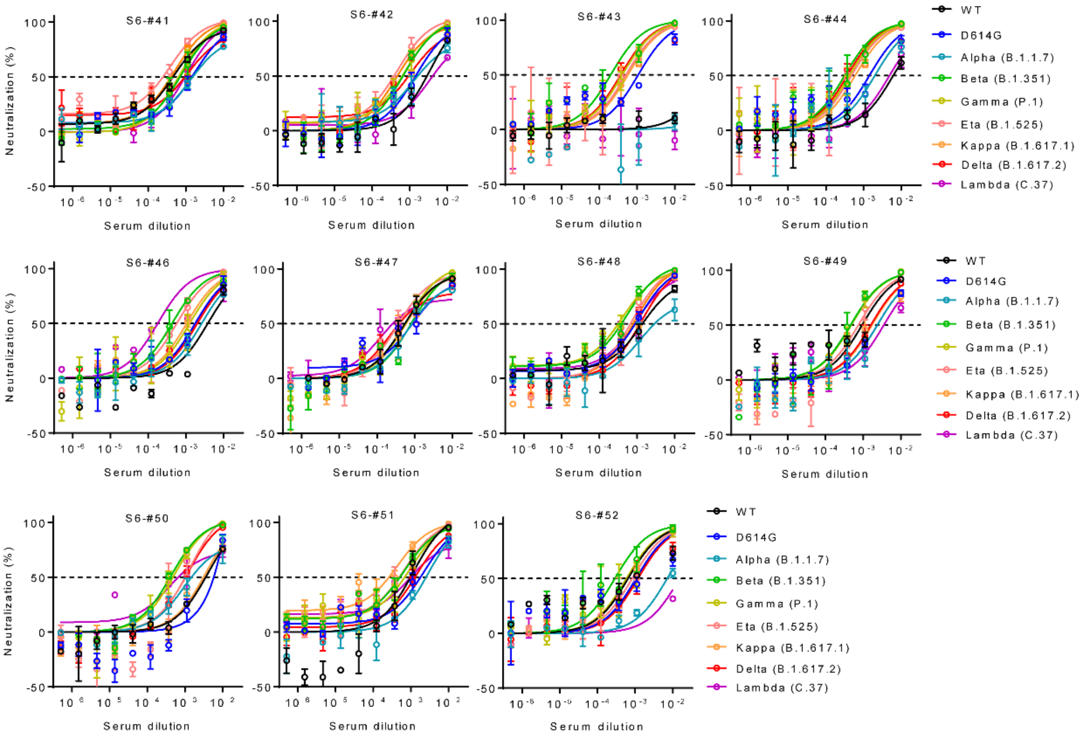


**Figure S5.** Neutralization of SARS-CoV-2 variants pseudoviruses in sera of mice injected $S_6$ subunit vaccine. Related to **Figure 3f**.

**Table S1.** List of monoclonal antibodies in this study.

| Serial number | Cat# | Property - Source | Company |
|---|---|---|---|
| #1 | 40590-D001 | monoclonal anti-human | SinoBiological |
| #2 | 40150-D002 | monoclonal anti-human | SinoBiological |
| #3 | 40150-D003 | monoclonal anti-human | SinoBiological |
| #4 | 40150-D004 | monoclonal anti-human | SinoBiological |
| #5 | 40150-D005 | monoclonal anti-human | SinoBiological |
| #6 | 40150-D006 | monoclonal anti-human | SinoBiological |
| #7 | 40592-MM57 | monoclonal anti-mouse | SinoBiological |
| #8 | 40150-R007 | monoclonal anti-rabbit | SinoBiological |
| #9 | 40591-MM42 | monoclonal anti-mouse | SinoBiological |
| #10 | 40150-D001 | monoclonal anti-human | SinoBiological |
| #11 | A02057-100 | monoclonal anti-mouse | genscript |
| #12 | 40592-R001 | monoclonal anti-rabbit | SinoBiological |
| #13 | 40591-MM43 | monoclonal anti-mouse | SinoBiological |
| #14 | GTX632604 | monoclonal anti-mouse | GeneTex |
| #15 | GTX635671 | monoclonal anti-rabbit | GeneTex |
| #16 | GTX635656 | monoclonal anti-rabbit | GeneTex |
| #17 | GTX635654 | monoclonal anti-rabbit | GeneTex |
| #18 | GTX635672 | monoclonal anti-rabbit | GeneTex |
| #19 | GTX01555 | monoclonal anti-human | GeneTex |
| #20 | A02109-100 | monoclonal anti-human | genscript |
| #21 | A02110-100 | monoclonal anti-human | genscript |
| #22 | A02051-100 | monoclonal anti-rabbit | genscript |
| #23 | A02052-100 | monoclonal anti-rabbit | genscript |
| #24 | A02053-100 | monoclonal anti-rabbit | genscript |
| #25 | A02054-100 | monoclonal anti-rabbit | genscript |
| #26 | A02055-100 | monoclonal anti-mouse | genscript |
| #27 | A02056-100 | monoclonal anti-mouse | genscript |

**Table S2.** Frequency of single mutations

| **Mutants** | Frequency | **Mutants** | Frequency |
|---|---|---|---|
| L18F | 1.5 | L452R | 6.6 |
| T20N | 0 | **E484K** | 7.4 |
| P26S | 0.7 | **N501Y** | 18.4 |
| **del69-70** | 59.6 | A570D | 16.9 |
| D80A | 3.7 | **D614G** | 91.2 |
| D138Y | 1.5 | H655Y | 1.5 |
| **del144** | 33.8 | Q677H | 5.1 |
| R190S | 0 | **P681H** | 18.4 |
| D215G | 3.7 | A701V | 3.7 |
| A222V | 0.7 | T716I | 16.2 |
| del242-244 | 6.6 | S982A | 16.2 |
| R246I | 0 | T1027I | 0 |
| K417N/T | 2.2/0 | D1118H | 16.9 |
